# Starvation Induces Diverse Autophagic Response in Cardiomyocytes and Noncardiomyocytes

**DOI:** 10.1101/2022.06.10.495581

**Authors:** Xi-Biao He, Haozhi Huang, Yi Wu, Fang Guo

**Affiliations:** Laboratory of Stem Cell Biology and Epigenetics, School of Basic Medical Sciences, Shanghai University of Medicine and Health Sciences, Shanghai 201318, China; Department of Orthopaedic Surgery, Shanghai Tenth People’s Hospital Affiliated to Tongji University, Shanghai 200072, China

**Keywords:** cardiomyocyte, noncardiomyocyte, autophagy, starvation, rapamycin

## Abstract

Cardiac autophagy serves as a potential mechanism to be genetically, epigenetically and pharmaceutically manipulated to prevent or alleviate pathological conditions in heart diseases. One of the complexities of this manipulation comes from the cellular heterogeneity of the heart. In this study, we assessed the dynamics of nutrient deprivation-induced autophagy activation in four major cardiac cell types in vitro and in vivo and found a major difference between cardiomyocyte and other three types of noncardiomyocyte. In comparison to the constant increase and maintenance of autophagic activity in noncardiomyocytes, cardiomyocyte undergoes rapid gain and loss of autophagic activity. This difference contributes to at least two aspects of cardiomyocyte pathophysiology: high vulnerability to nutrient deprivation-induced cell death and unresponsiveness to pharmaceutical regulation of autophagy. Our present findings support a cell type-specific targeting of autophagy for therapeutic intervention of cardiac diseases.

## INTRODUCTION

Heart is composed of a variety of cell types, each of which possess unique molecular and cellular characteristics determining their physiological functions and pathological responses. Several recent studies using single-cell sequencing methodologies have depicted cell type-specific dynamics of gene expression profile, intercellular communication and pathophysiology in heart development and disease [17, 18, 21], highlighting the importance of systematic and comparative analysis of all the cardiac cell types for a comprehensive understanding of heart behaviors. In particular, stage-specific crosstalk between cardiomyocytes and noncardiomyocytes have been extensively implicated in the progress of cardiac diseases such as hypertrophy and heart failure (HF) [17, 18], further emphasizing the necessity of taking these factors into consideration for therapeutical interventions.

Macroautophagy/autophagy is an evolution-conserved intracellular catabolic process utilized by all mammalian cells to maintain proteostasis and defend against intrinsic and extrinsic stresses. In heart, all cardiac cells are equipped with proper autophagic activity (basal/constitutive autophagy) and able to undergo autophagy activation (adaptive autophagy) in response to a wide range of cardiac stresses, including misfolded protein aggregation, aging and starvation [9]. A large number of studies have shown induction of autophagy in cardiomyocytes in response to acute and chronic myocardial infarction (MI), ischemia/reperfusion, hypertrophy, cardiomyopathy and HF, although it remains controversial whether the outcome is definitely beneficial or detrimental [9]. However, few have set noncardiomyocytes as parallel goals of research to avoid the overlook of their roles in the pathology of cardiomyocytes and whole heart. In addition, regulation of autophagy through pharmaceutical or life style interventions has been proposed as a novel preventive or therapeutic strategy against MI-related cardiac impairments, which again mainly focuses on the autophagic activity of surviving cardiomyocytes [2, 6, 23]. How noncardiomyocytes respond to autophagy-related stimuli and to what extent do they differ from cardiomyocyte, remain largely unexplored.

Starvation is a potent inducer of autophagy. Cardiac starvation caused by nutrient deprivation is a common pathology in human heart diseases such as MI, late-stage HF, cardiac remodeling and pathological aging [1, 9]. It is widely accepted that one of the most affected cardiac cell types in these diseases is cardiomyocyte, malfunction and loss of which are directly associated with a variety of cardiac dysfunction. One fact that is often less emphasized, however, is that noncardiomyocytes which generally account for more than 70% of the total amount of cells in mouse and human heart [16], are exposed to similar intensity of stress as cardiomyocyte and undergo even more severe cell damage [3]. For instance, the majority of apoptotic cells in monkey and human hearts suffered from MI and HF have been proven to be noncardiomyocytes, including macrophage, neutrophils, fibroblasts and endothelial cells [15]. Given their diverse developmental origins and functions [11], a large variety of adaptive behaviors are expected from each type of noncardiomyocytes in response to cardiac starvation, creating a highly complex cellular niche together with cardiomyocyte. The discrepancy of how cardiomyocyte and noncardiomyocytes survive cardiac stresses might greatly affect their pathogenesis and the strategy of corresponding therapeutic interventions.

Here, by monitoring the changes of autophagy markers using several autophagy assessment methods, we have provided a dynamic pattern of autophagy activation upon starvation in a primary heterogeneous cell culture of four major cardiac cell types and confirmed the consistency with that in starved neonatal heart. A key difference of autophagy characteristics between cardiomyocyte and other three noncardiomyocytes is revealed, which to a large extent distinguishes how they survive and respond to pharmaceutical autophagy regulators during starvation.

## Materials and Methods

### Ethics

Neonatal ICR mouse pups were supplied by Shanghai Jiesijie Experimental Animal Co. All animal experiments were approved by the animal ethics committee of Shanghai University of Medicine & Health Sciences and have been performed in accordance with the ethical standards laid down in the 1964 Declaration of Helsinki and its later amendments.

### Cell culture and starvation induction

Three cell culture containing different composition of cardiac cell types were performed as previously described [5]. Briefly, after decapitation, the ventricles of postnatal day 1 mouse pups were quickly dissected under a microscope. For heterogeneous cell culture containing both cardiomyocytes and noncardiomyocytes, tissues were enzymatically digested and mechanically pipetted into single cells and plated onto laminin-coated culture coverslips, plates or dishes. For homogeneous cardiomyocyte culture, a cardiomyocyte growth supplement from Pierce primary cardiomyocyte isolation kit (Thermo Scientific) was added to reduce noncardiomyocyte growth during cell culture periods. For noncardiomyocyte culture, heterogeneous cell culture was passaged once and mechanical pipetted to eliminate all cardiomyocytes. The remaining noncardiomyocytes were re-plated. All cells were applied to experiments 5 days after plating. Cells were grown in Dulbecco’s modified Eagle medium supplemented with 10% fetal bovine serum and 1% Penecillin/Streptomycin and incubated in 5% CO_2_, 37□ incubators. To induce starvation in vitro, cells were washed by Hank’s balanced salt solution (HBSS) twice to eliminate medium residue and incubated in HBSS for various periods of time. For starvation induction in vivo, postnatal day 1 mouse pups were starved by milk depletion for various periods of time and sacrificed by decapitation.

### Fluorescence immunostaining analysis

After decapitation, heart tissues were washed with cold HBSS twice, immediately fixed in cold 4% paraformaldehyde for 2 hours and dehydrated in 30% sucrose. Tissues were then stored in -80□ until they were cryosectioned into 20 μm thickness (Leica CM1950). Cells were fixed with 4% paraformaldehyde for 20 minutes followed by phosphate buffered saline wash three times. For immunostaining, fixed cells or tissue slices were permeabilized and blocked in phosphate buffered saline with 0.3% Triton-X100 and 1% bovine serum albumin for 40 minutes, then incubated with primary antibodies diluted with blocking solution at 4□ overnight. Alexa Fluor series of second antibodies (Thermo Scientific) were applied accordingly for one hour at room temperature. Cells were finally mounted in 4’,6-diamidino-2-phenylindole (DAPI) and examined using fluorescence microscope (Leica DMi8). The first antibodies include rabbit mouse anti-sarcomeric α-actinin (SAA; Abcam), rabbit anti-discoidin domain-containing receptor 2 (DDR2; Immunoway), mouse anti-smooth muscle actin (SMA; Cell Signaling Technology), Lectin from tomato, FITC conjugate (Sigma), rabbit anti-microtubule-associated protein 1 light chain 3 (LC3) and mouse anti-LC3 (both from MBL).

### Western blotting analysis

Cells were lysed for 30 minutes in lysis buffer with protease inhibitor cocktails (Sigma). 3 μg of proteins were separated by SDS-polyacrylamide gel electrophoresis and transferred to PVDF membrane. The membrane was blocked with 5% skim milk (Cell Signaling Technology) for 1 hour at room temperature, incubated with first antibodies including mouse anti-LC3, mouse anti-β-actin and rabbit anti-p62 (both from Cell Signaling Technology) overnight at 4 °C. The membranes were washed three times with Tris-buffered saline and Tween-20 followed by incubation with the peroxidase-conjugated anti-mouse or anti-rabbit secondary antibodies (both from Millipore) for 1 h at room temperature.

### Cell counting and statistics

Immunoreactive or DAPI-stained cells were counted in at least 10 random regions of each culture coverslip using an eyepiece grid at a magnification of 50 to 400X. Data are expressed as mean ±\ S.E.M. of three independent cultures. Statistical comparisons were made using Student’s t-test or one-way ANOVA with Tukey’s post hoc analysis (Graphpad Prism).

## RESULTS

### Autophagic activity is rapidly declined in nutrient-deprived cardiomyocytes

To determine the kinetics of starvation-induced autophagy in different cardiac cell types in vitro, we recruited a previously well-defined primary heterogeneous cell culture composed of four major cardiac cell types, including SAA+ cardiomyocytes, DDR2+ cardiac fibroblasts, SMA+ vascular smooth muscle cells and Lectin+ endothelial cells [5]. Autophagic activity was quantified by the number of LC3 puncta in each cell expressing corresponding cell type markers. Within 3 hours of HBSS-induced starvation, a rapid increase of LC3 puncta number was observed in 10 min, dramatically decreased in 30 min, and maintained at negligible level for the rest of time (Fig. 1A). Of note, a considerable amount of LC3 puncta existed in nutrient-rich cardiomyocytes, which could be further agonized by treatment of Bafilomycin A1 (BAF; data not shown), suggesting that the autophagic flux is intact and a basal autophagy is needed for cardiomyocyte physiology.

**Figure 1.**
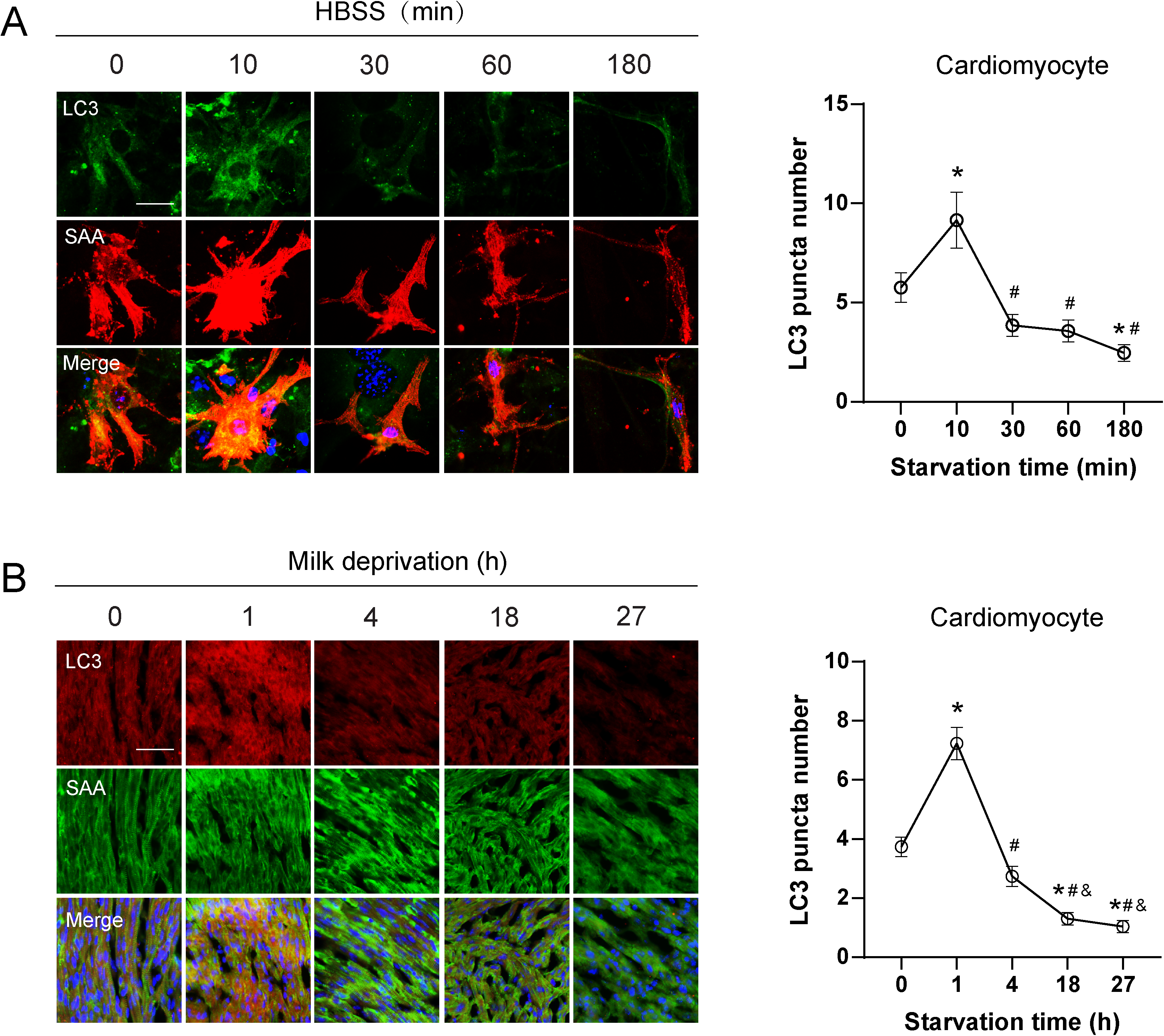
Dynamics of autophagic activity in cardiomyocytes in response to starvation. (A) Representative immunofluorescence images and quantification of LC3 puncta in sarcomeric α-actinin (SAA)-marked cardiomyocytes in heterogenous cardiac cell culture. Starvation was induced by incubating cells in Hank’s balanced salt solution (HBSS) for 10 to 180 min. Scale bar represents 20 μm. Data represent mean ± S.E.M. \**P* < 0.05 vs 0 min; #*P* < 0.0001 vs 10 min; One-way ANOVA with Tukey’s post hoc test. (B) Representative images and quantification of LC3 puncta in SAA+ cardiomyocytes residing in the left ventricle myocardium of mouse neonates starved for 1 to 27 hours. Scale bar represents 50 μm. A minimum of 50 cells per sample was counted from three independent experiments. Data represent mean ± S.E.M. \**P* < 0.0001 vs 0 h; #*P* < 0.0001 vs 1 h; & *P* < 0.05 vs 4 h; One-way ANOVA with Tukey’s post hoc test.

To confirm our in vitro data, the dynamics of LC3 expression was assessed in cardiomyocytes residing in the left ventricular myocardium of mouse neonates starved for 1, 4, 18 and 27 hours. Co-labeling SAA+ cardiomyocytes with LC3 by immunostaining analysis demonstrated that LC3 expression was maintained at basal level in SAA+ cardiomyocytes and immediately increased within 1 hour of milk depletion-induced starvation, followed by a gradual decrease within 4 hours (Fig. 1B). Thereafter, the LC3 expression reached undetectable level, suggesting a severe loss of autophagic activity after prolonged starvation. This finding is in line with previous study monitoring autophagy dynamics using transgenic GFP-LC3 mice [8]. Taken together, these results unravel a rapid gain and loss dynamics of autophagy in cardiomyocytes upon starvation.

### Autophagic activity is slowly increased in nutrient-deprived noncardiomyocytes

To determine the autophagy dynamics of other three cardiac cell types in parallel with cardiomyocyte, same experimental procedure as that of cardiomyocyte was applied to these cells in vitro. In contrast to cardiomyocyte, all three types of noncardiomyocyte exhibited low basal autophagy, as evidenced by negligible LC3 puncta in nutrient-rich condition. Treatment of BAF further induced an increase of LC3 puncta, confirming the autophagic flux integrity in these cells (data not shown). Within 10 min of starvation, the LC3 level in DDR2+ cardiac fibroblasts dramatically increased, and was maintained so for 3 hours (Fig. 2A). As to SMA+ vascular smooth muscle cells, the LC3 level peaked within 1 hour of starvation and gradually decreased afterwards (Fig. 2B). Similarly, Lectin+ endothelial cells also underwent gradually increase of LC3 within 1 hour of starvation, yet with a much milder extent. Diversely, the highest level of LC3 in endothelial cells appeared at the time point of 3 hours (Fig. 2C), suggesting a boost of autophagic activity after a long-term period of starvation. Nevertheless, these three cell types of noncardiomyocyte exhibited similar trend of autophagy activation in response to short-term, i.e., 1 hour of starvation, which is remarkably distinguished from that of cardiomyocyte. In consistent with in vitro observations, starved vascular endothelial cells in the left ventricular myocardium of mouse neonates exhibited similar dynamics of autophagic activity, as evidenced by LC3 expression in Lectin+ cells in vivo (Fig. 2D).

**Figure 2.**
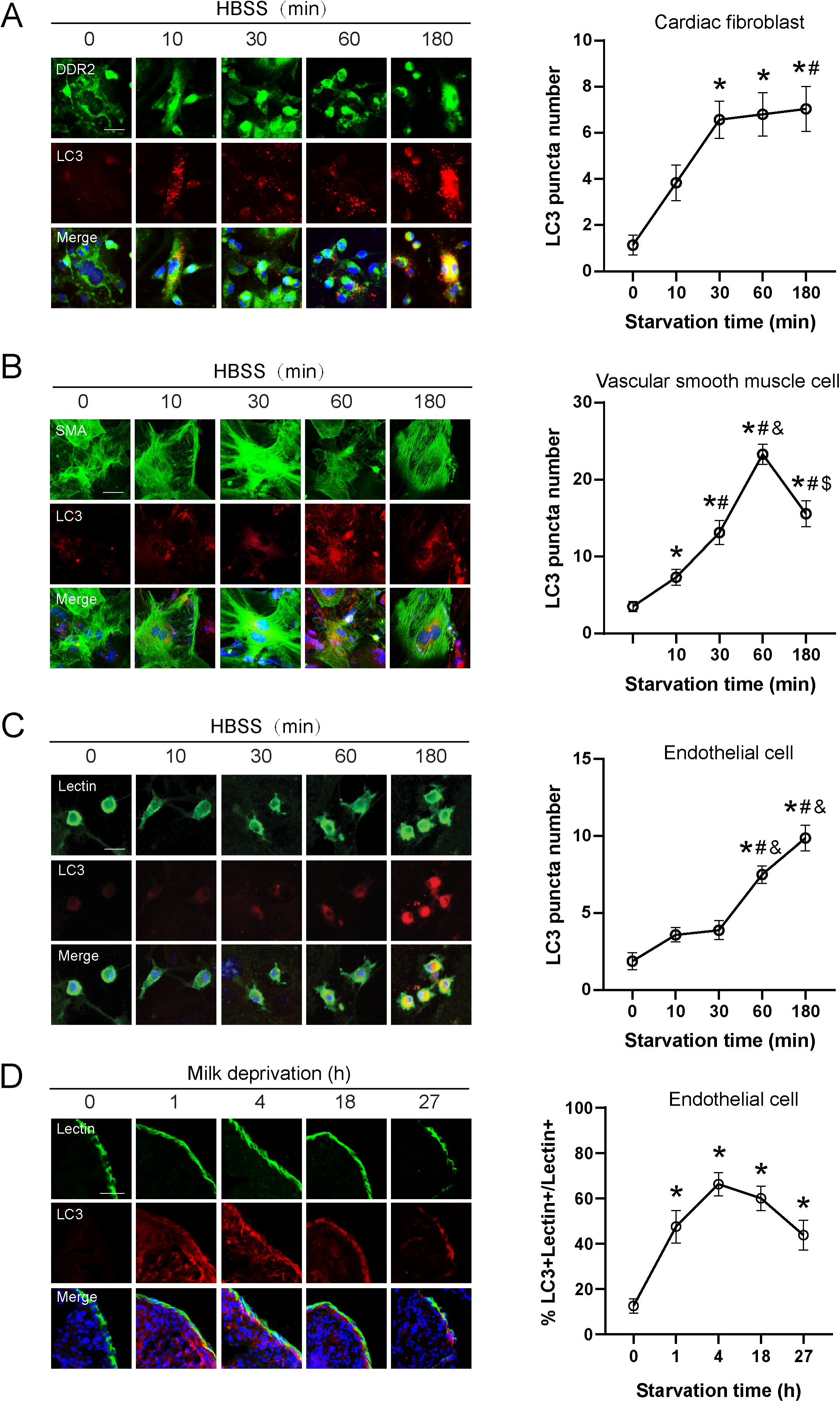
Dynamics of autophagic activity in noncardiomyocytes in response to starvation. (A-C) Representative immunofluorescence images and quantification of LC3 puncta in discoidin domain-containing receptor 2 (DDR2)-marked cardiac fibroblasts, smooth muscle actin (SMA)-marked vascular smooth muscle cells and Lectin-marked endothelial cells in heterogenous cardiac cell culture. Starvation was induced by incubating cells in Hank’s balanced salt solution (HBSS) for 10 to 180 min. Scale bar represents 20 μm. Data represent mean ± S.E.M. \**P* < 0.0001 vs 0 min; #*P* < 0.05 vs 10 min; &*P* < 0.0001 vs 30 min; $*P* < 0.05 vs 60 min; One-way ANOVA with Tukey’s post hoc test. (D) Representative images and quantification of Lectin+ endothelial cells expressing Lectin residing in the left ventricle myocardium of mouse neonates starved for 1 to 27 hours. Scale bar represents 50 μm. A minimum of 50 cells per sample was counted from three independent experiments. Data represent mean ± S.E.M. \**P* < 0.05 vs 0 h, One-way ANOVA with Tukey’s post hoc test.

### Opposite responsiveness to rapamycin between cardiomyocyte and noncardiomyocytes

The differential autophagic dynamics in response to starvation between cardiomyocyte and noncardiomyocytes prompted us to hypothesize that they would also respond differentially to autophagy regulation signals. Rapamycin serves as a widely-studied agonist of starvation-related autophagy. To test our hypothesis, we examined how autophagy was altered by rapamycin in starved cardiomyocytes and noncardiomyocytes by analyzing the expression dynamics of LC3 and p62, a common starvation-related autophagy receptor by Western blotting experiments. To enable a parallel comparison without cross contamination, the heterogenous cell culture was optimized to generate a homogeneous cardiomyocyte culture and a noncardiomyocyte culture by previously established methods in our lab (Fig. 3A) [5].

**Figure 3.**
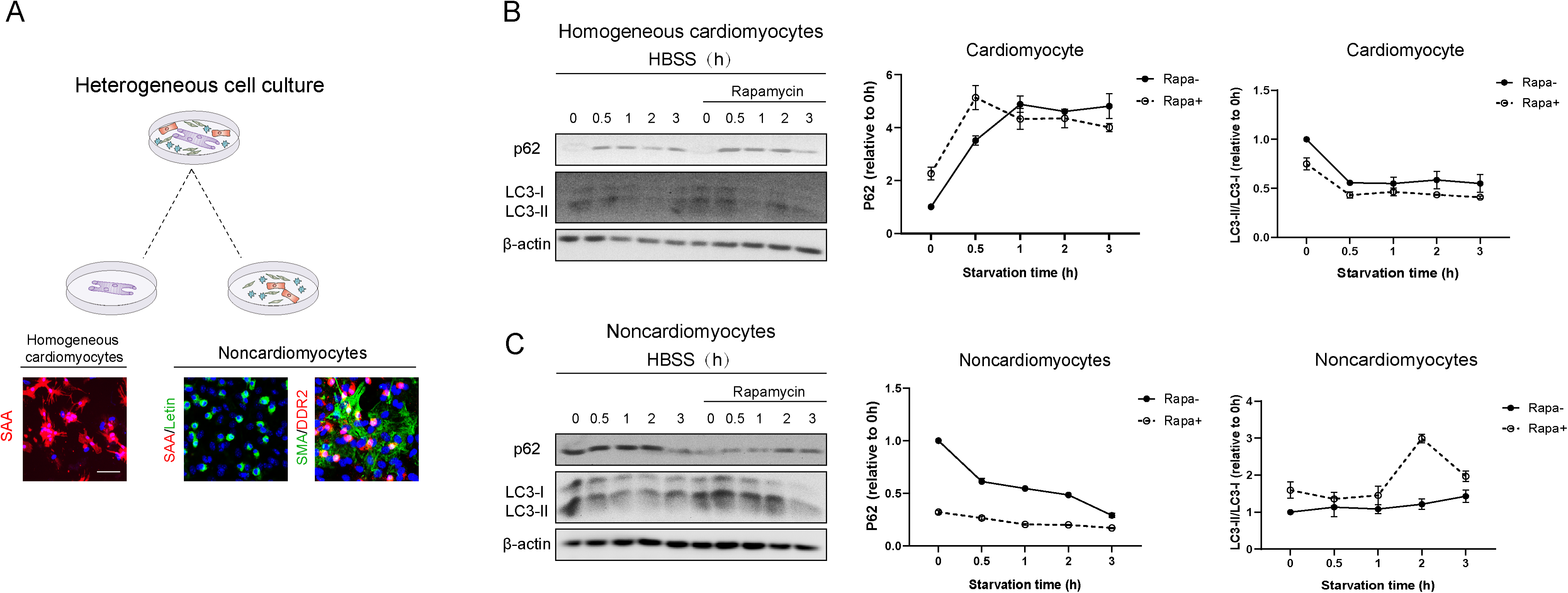
Dynamics of autophagic activity in response to rapamycin in starved cardiomyocytes and noncardiomyocytes. (A) Schematics depicting the experimental procedure of homogeneous cardiomyocyte culture and noncardiomyocyte culture derived from heterogenous cardiac cell culture. The composition of each culture is validated by representative immunofluorescence images marking cardiomyocytes expressing sarcomeric α-actinin (SAA), cardiac fibroblasts expressing discoidin domain-containing receptor 2 (DDR2), vascular smooth muscle cells expressing smooth muscle actin (SMA) and endothelial cells expressing Lectin. Scale bar represents 20 μm. (B and C) Representative images and densitometry of Western blotting analysis on autophagy markers LC3 and p62 from samples collected from pure cardiomyocyte culture and noncardiomyocyte culture, respectively. Cells were starved in Hank’s balanced salt solution (HBSS) for 30-180 min. Rapamycin was pre-treated 10 min before starvation. 0 min with rapamycin indicates 180 min incubation in cell culture media with rapamycin. Densitometry of each band was measured by ImageJ software and normalized with β-actin from two independent experiments.

Consistent with immunostaining results in Fig. 1, the level of LC3-II:LC3-I ratio in cardiomyocytes was decreased within 3 hours of starvation, whereas that of p62 was inversely increased, confirming a loss of autophagic activity in cardiomyocytes upon starvation. Rapamycin treatment during the whole starvation period neither increased LC3 ratio nor decreased p62 expression, indicating a failed rescue of the autophagic activity (Fig. 3B). Importantly, addition of rapamycin in the absence of starvation stimulation did increase the LC3 ratio and decrease p62 expression in comparison to no rapamycin treatment, confirming the efficiency of rapamycin on activating basal autophagy in cardiomyocytes.

By contrast, in noncardiomyocyte culture consisting of the other three cardiac cell types, the levels of LC3-II:LC3-I ratio and p62 were largely unaltered during hours of starvation, demonstrating a stable maintenance of autophagic activity (Fig. 3C). The higher intensity of LC3 ratio without starvation might be derived from the heterogeneity nature of cells. Importantly, rapamycin treatment remarkably induced an increase of LC3 ratio and a decrease of p62 expression from 0.5 to 2 hours of starvation, suggesting a substantial enhancement of autophagic activity. These results not only confirmed the distinct autophagy dynamics between cardiomyocytes and noncardiomyocytes upon starvation, but also indicated a critical difference in their autophagic responsiveness to pharmaceutical autophagy agonists.

### Starved cardiomyocytes are insensitive to pharmaceutical autophagy regulation

The rapid loss of autophagic activity in starved cardiomyocytes appears to suggest that these cells are more vulnerable to nutrient deprivation than noncardiomyocytes. Indeed, 2 hours of starvation already reduced about half of the cardiomyocytes in heterogeneous cell culture, whereas the viability of noncardiomyocytes was not significantly affected (Fig. 4A). We asked whether positive or negative regulation of autophagy would affect the viability of starved cardiomyocytes. As expected, addition of rapamycin did not rescue the loss of cardiomyocytes. Interestingly, inhibition of autophagy by 3-methyadenine (3-MA) or BAF together with starvation did not further reduce the cell number of cardiomyocytes (Fig. 4B). This unexpected result suggested an insensitivity of cardiomyocyte to exogeneous regulation of autophagy. In summary, our findings suggest that the survival of cardiomyocytes under nutrient-deprived condition is determined by their unique characteristics of autophagy, which is largely distinguished from noncardiomyocytes.

**Figure 4.**
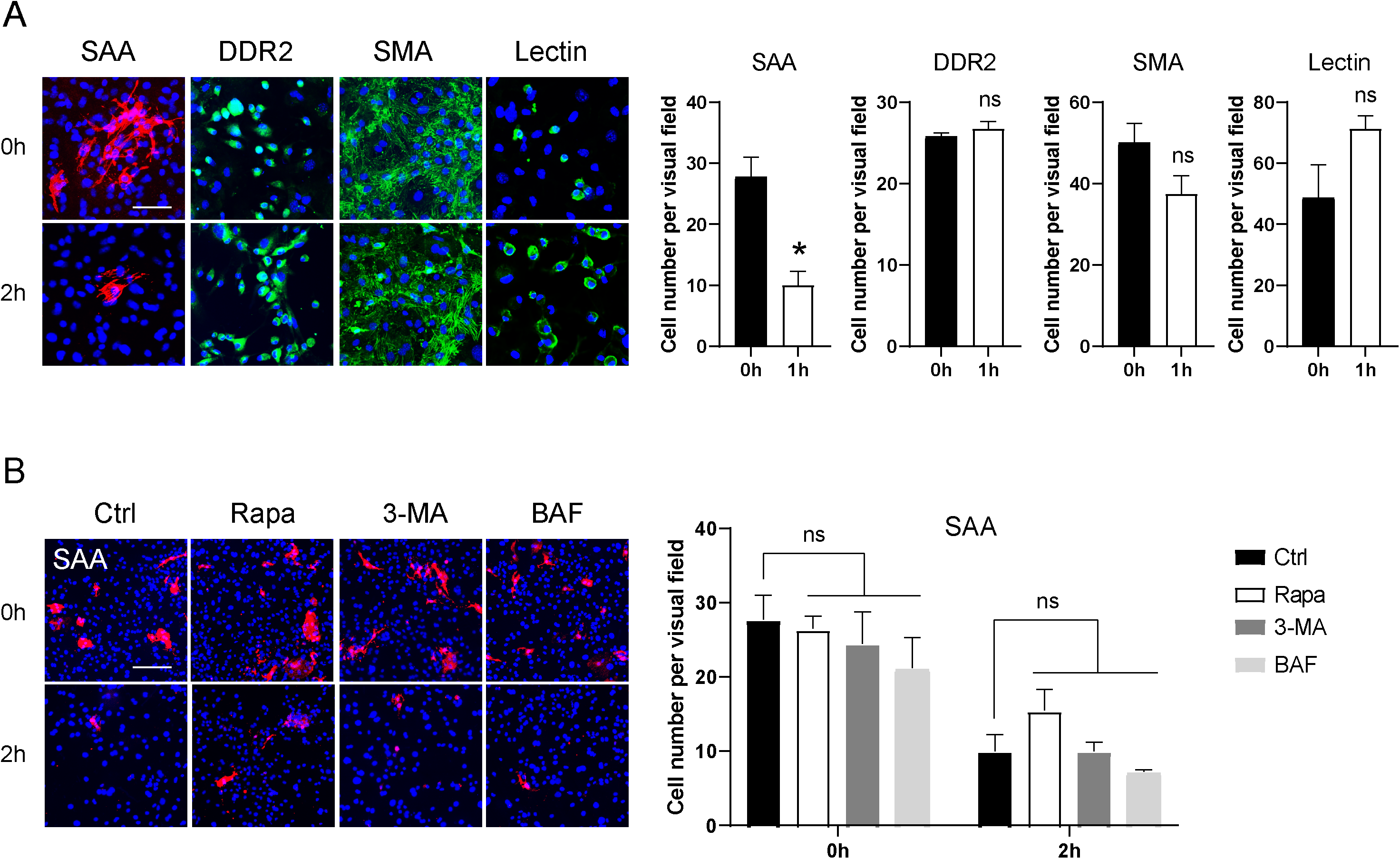
Survival of cardiomyocytes and noncardiomyocytes in response to starvation and pharmaceutical autophagy regulation. (A) Representative immunofluorescence images and quantification of cell number of sarcomeric α-actinin (SAA)-marked cardiomyocytes, discoidin domain-containing receptor 2 (DDR2)-marked cardiac fibroblasts, smooth muscle actin (SMA)-marked vascular smooth muscle cells and Lectin-marked endothelial cells before and after 2 hours of Hank’s balanced salt solution-induced starvation. (B) Representative immunofluorescence images and quantification of cell number of SAA-marked cardiomyocytes before and after 1 hour of Hank’s balanced salt solution-induced starvation with or without the presence of positive autophagy regulator rapamycin (Rapa), negative autophagy regulator 3-methyadenine (3-MA) or Bafilomycin A1 (BAF). All drugs were 10 min pre-treated before starvation. Cell numbers were counted in 10 random areas of each culture coverslip using an eyepiece grid at a magnification ×100. Scale bar represents 20 μm. n = 3 independent experiments. Data represent mean ± S.E.M. \**P* < 0.05, ns not significant; Student’s t-test and Two-way ANOVA with Dunnett’s post hoc test.

## DISCUSSION

The main finding of this study is that the characteristics of autophagy distinguishes cardiomyocyte and noncardiomyocytes in starved heart. In contrast to the rapid loss of autophagic activity in cardiomyocyte, noncardiomyocytes, including vascular smooth muscle cell, endothelial cell and cardiac fibroblast, possess extended duration of autophagic activation in response to starvation. In addition, the cardiomyocyte-specific loss of autophagic activity at least partially contributes to their low viability and unresponsiveness to pharmaceutical autophagy regulators under cardiac starvation, a state that is largely reminiscent of limited nutrient and oxygen supply to myocardium in ischemia (Figure 5). We propose that the cardiac cell type-specific difference of autophagic activation in response to starvation in vitro might provide evidence for the understanding of their behavior under ischemia in vivo.

**Figure 5.**
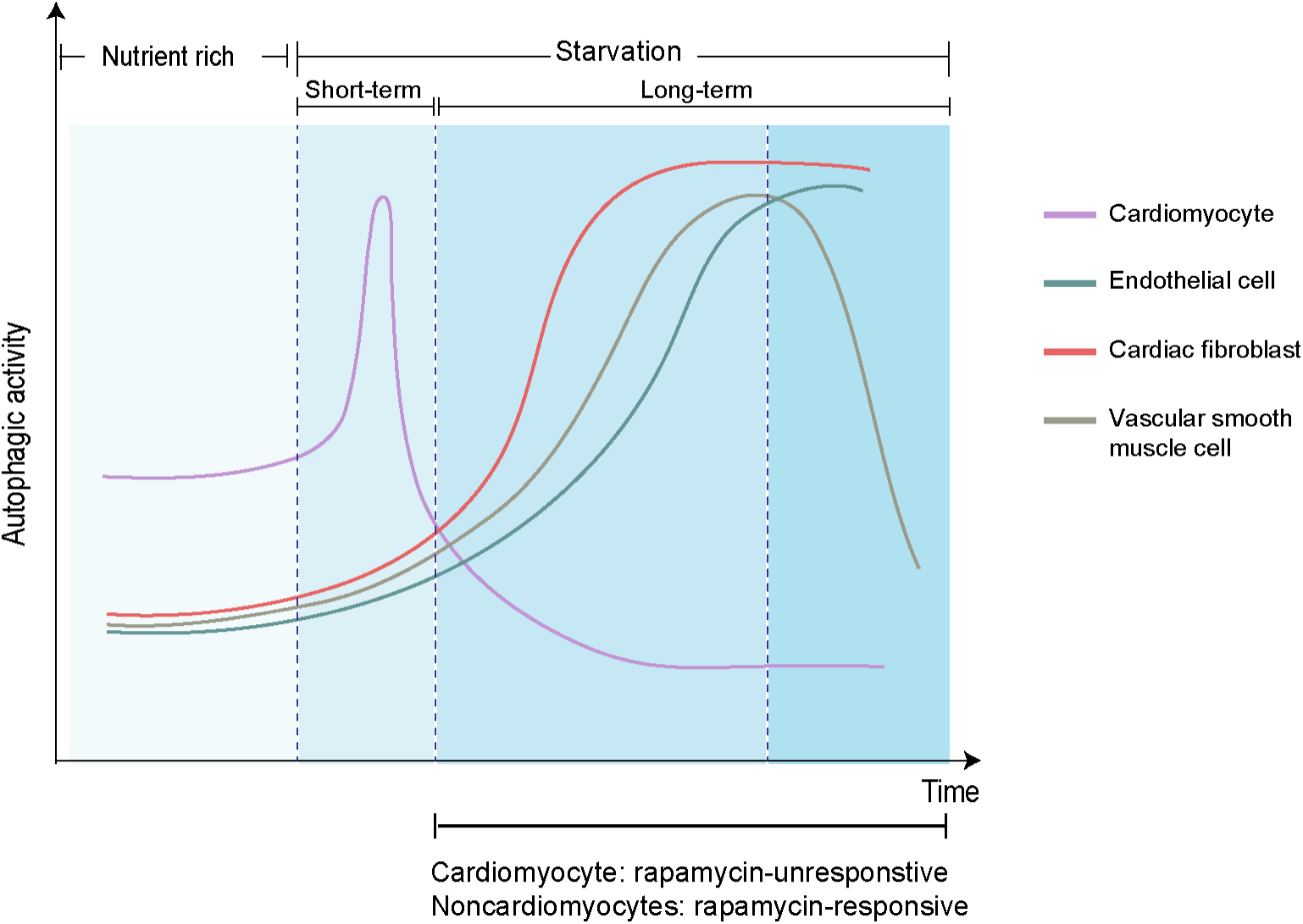
Schematics summarizing the cardiac cell type-specific dynamics of autophagic activity in response to starvation in vitro. In nutrient-rich condition, cardiomyocytes exhibit higher level of basal autophagy than noncardiomyocytes such as cardiac fibroblasts, vascular smooth muscle cells and endothelial cells. Upon starvation, the autophagic activity in cardiomyocytes is gained and lost within very short period of time, whereas that in all three noncardiomyocytes is gradually increased. Thereafter, the autophagic activity in noncardiomyocytes is further increased and maintained at high level for a much longer period of time, with concomitant negligible low level of autophagic activity in cardiomyocytes. In this period, cardiomyocytes exhibit unresponsiveness to pharmaceutical autophagy regulation and susceptibility to cell death.

Autophagy capacity is one of the most fundamental characteristics of eukaryotic cells that determines a wide range of biological functions including energy catabolism, protein homeostasis and stress response. Both basal autophagy and adaptive autophagy are essential for the normal function of heart in health and disease [8, 13, 14, 19, 20]. Our results showed that both basal autophagy and adaptive autophagy in response to starvation varied amongst different cardiac cell types. In nutrient-rich condition, plenty of LC3 puncta were already enriched in cardiomyocyte, whereas those in three types of noncardiomyocytes were barely detected (Fig. 1A and 2A-C). This reflects a cardiomyocyte-specific high turnover of intracellular components thereby emphasizing the importance of basal autophagy to protein homeostasis in cardiomyocyte. In line with this conclusion, cardiomyocyte-specific Atg5 knockout in transgenic mice causes severe cardiomyopathy and protein aggregation [14]. Upon starvation, however, we found that the autophagic activity in cardiomyocyte was dramatically lost within half hour, accompanied by a rapid aggregation of p62 (Figure. 3B), suggesting an attenuated degradation of autophagic cargo. By contrast, noncardiomyocytes were able to activate adaptive autophagy by gradually generating more LC3 autophagosomes and maintaining the autophagic activity within 3 hours of starvation (Fig. 3C). As a consequence, p62 expression was maintained in a stable level and could be further reduced by rapamycin. Indeed, p62 has been shown to be one of the most efficiently degraded autophagy substrates in response to starvation [12]. The inverse association of p62 expression and the conversion of LC3-I to LC3-II validates the bona fide autophagic activity in our work.

It is apparent that the autophagy dynamics during starvation alter in a cell type- and time-specific manner. The cardiomyocyte-specific up- and down-regulation of LC3 within 10 min of starvation in vitro and 1 hour in vivo indicates that starvation induces a rapid gain and loss of autophagic activity rather than an impairment of autophagy mechanism. By contrast, all three types of noncardiomyocytes were able to elevate and maintain their autophagic activity at high level within 2 hours of starvation in vitro, suggesting great varieties in their autophagic response with cardiomyocyte. Researches on cell type diversity of autophagy is surprisingly rare. However, accumulating evidence exist from recent single-cell sequencing studies showing that the gene expression profile of a number of autophagy-related genes are largely differed in different cardiac cell types [17, 18, 21]. Screening of the most divergent genes and validating their roles in cell type-specific autophagic response might help unravel the mechanisms underlying this diversity in future study.

Whether insufficient autophagy activity is in a causal relationship or associated with cardiomyocyte cell death remains elusive. Our results support the idea that the feature of poor autophagy plasticity in cardiomyocyte might at least partially contribute to the deterioration of its survival. Although about half of the cardiomyocytes were already lost in response to 2 hours of starvation in vitro (Fig. 4A), no obvious cardiomyocyte cell death was observed in starved neonatal heart (data not shown). The discrepancy might be explained by the fact that cardiomyocytes in heart are much well supported by the integral circulation system in vivo. In support, previous study has shown that starvation for 3 days in adult heart does not induce cardiomyocyte apoptosis, but the ATP content in cardiomyocytes is further decreased by BAF-induced inhibition of autophagy and cardiac dysfunction is observed [7]. Interestingly, in the same study, TUNEL+ apoptosis was induced by BAF in noncardiomyocytes, suggesting that the survival of noncardiomyocytes is more sensitive to autophagy inhibition. It is also reported that the majority of cell death in HF occurs in noncardiomyocytes [15]. Our finding demonstrating that the cardiomyocyte-specific low responsiveness to negative autophagy regulators provides a novel insight on this issue.

The pathophysiological consequences resulted from the diverse patterns of autophagic activity between cardiomyocyte and noncardiomyocytes remain unclear. A number of evidence have demonstrated the involvement of autophagy in different noncardiomyocytes in the context of disease. For instance, MI induces impairment of autophagic flux in cardiac fibroblasts and results in cardiac remodeling and ventricular dysfunction after MI [22]. In vascular endothelial cells and smooth muscle cells, hypoxia induces autophagy activation in these cells through elevation of angiogenic factor AGGF1, lack of which reduces protective angiogenesis in MI [10]. These studies emphasize the importance of maintaining or increasing autophagic activity in noncardiomyocytes for the prevention of cardiac pathologies. Based on our results, it appears intriguing to further assess the effects of autophagic activity-depleted cardiomyocytes on neighboring noncardiomyocytes, considering that the preceding deterioration of cardiomyocytes due to loss of autophagic activity might result in a secondary, or even amplifying detrimental effect on surrounding noncardiomyocytes. From this point of view, It is reported that even very low level of cardiomyocyte apoptosis is sufficient to cause HF [24], further highlighting potential intercellular communications between dying cardiomyocyte and surviving noncardiomyocytes that might lead to sequential pathogenesis in different cardiac cell types.

Autophagy activation with pharmaceutical positive regulators such as rapamycin serves as a potential therapy for heart diseases. The unresponsiveness of starved cardiomyocytes to rapamycin observed in our study might, in theory, suggest that the beneficial effects of rapamycin come from alternative cell sources such as noncardiomyocytes. This idea is in line with the emerging awareness and calling for cell-type-targeted intervention of cardiac diseases. Noncardiomyocytes including endothelial cells and smooth muscle cells have been shown to be actively involved in the performance of cardiomyocyte and cardiac function [4]. The mechanisms by which manipulating noncardiomyocyte autophagy benefits the progression of cardiac diseases deserve further investigation.

## ABBREVIATIONS

BAF: Bafilomycin A1
DAPI: 4′,6-diamidino-2-phenylindole
DDR2: Discoidin domain-containing receptor 2
HBSS: Hank’s balanced salt solution
HF: Heart failure
LC3: Microtubule-associated protein 1A/1B-light chain 3
MI: Myocardial infarction
PBS: Phosphate buffered saline
SAA: Sarcomeric α-actinin
SMA: Smooth muscle actin
3-MA: 3-methyadenine

## Acknowledgements

This work was supported by the National Natural Science Foundation of China (31701287 and 32100779).

## Data Availability Statement

The data that support the findings of this study are available from the corresponding author upon reasonable request.

